# Are mitochondrial DNA mutations under purifying selection in somatic tissues?

**DOI:** 10.1101/2025.10.26.684677

**Authors:** Melissa Franco, Jacob Bandell, Konstantin Popadin, Dori Woods, Konstantin Khrapko

## Abstract

The extent to which somatic mitochondrial DNA (mtDNA) mutations are subject to selection is a fundamental question relevant to development, mitochondrial disease, cancer, and aging. Recently a study from the Sudmant laboratory that used an advanced, high fidelity mutational analysis reported that somatic mutations in protein-coding genes exhibit signatures of negative selection. This report came as surprise as several other studies including those that used same technology reported either lack of selection or positive (destructive) selection on somatic mutations. We hypothesized that these discrepancies may stem, in part, from the inclusion of germline mutations in addition to somatic ones, which could bias selection analyses due to the high synonymity of the latter. To test this, we reanalyzed the Sudmant dataset by separating mutations into germline (defined as shared between related animals) and somatic (not shared between tissues of an animal). We then employed a cumulative curve approach to assess selection without bias. Our analysis reveals that, indeed, an apparent purifying selection signal is driven by an admixture of synonymous germline mutations and disappears upon their removal. The remaining somatic mutations for most part show overall dynamics consistent with neutral drift. However, mutations at higher mutant fractions show positive selection trend, most compatible with a low proportion of mutations experiencing positive selection. While we do not exclude rare or context-specific selection events, our results argue against pervasive somatic selection and highlight the importance of rigorous stratification when interpreting mtDNA mutational patterns.

## Introduction

Selection of mtDNA mutations in somatic tissues is pertinent to mtDNA-based disease, aging and cancer. A large number of studies in the past attempted to address these questions and results varied. In mtDNA-based diseases, it’s often observed that the fraction of mutations in the affected tissues increases with time. In cancer, perhaps the most reliable study (Yuan et al., 2020) has determined that while most mutations in cancers appear to be not under selection, some mutations (i.e., highly detrimental truncating mutations) may be either under negative selection in most cancers, but under positive selection in a few special cases (kidney, colorectal, thyroid). A classic example is the MELAS mutation (m.3243A>G), which appears to be under negative selection in blood but no selection in muscle (Grady et al., 2018). It’s typical to see an increase of mutational load with age, though in many cases it is difficult to determine whether the increase is due to positive selection of existing mutations or de novo mutational accumulation (Marcelino et al., 1998) succeeded with neutral clonal expansion of nascent mutations (which is not selection). In some cases positive intracellular selection has been explicitly demonstrated for deletions (Bua et al., 2006), (Nicholas et al., 2009). We ourselves have reported positive intracellular selection of specific somatic mtDNA mutations in single cells in 2002 (Nekhaeva et al., 2002). However, in our study of clonally expanded somatic mtDNA mutations in colonic crypts (Greaves et al., 2014), no selection was observed. In contrast, Stoneking reported positive somatic selection in heteroplasmic mutations, though their study dealt with very high fraction mutations, i.e. above 0.5% (Li et al., 2015). Very recently, support for positive ‘destructive’ somatic selection came from single cell analysis (Korotkevich et al., 2024). Interestingly, in the germline, both positive (Fleischmann et al., 2024), (Cote-LHeureux et al., 2023), (Zhang et al., 2021), and negative (Stewart et al., 2008) selection has been reported, making this issue very multifaceted.

Traditionally, the studies of somatic mutations were hampered by their low frequency and contamination with artifacts (Khrapko and Vijg, 2008). Recently, two studies used high-fidelity mutation analysis, ‘ds sequencing’, which greatly minimized these artifacts. Both studies addressed the somatic selection issue, and, intriguingly, gave different answers. Whereas the Kennedy lab (Sanchez-Contreras et al., 2023) and a previous similar but much smaller study (Arbeithuber et al., 2020) found no evidence for consistent selection, the Sudmant lab (Serrano et al., 2024) reported ‘signatures of negative selection’ and concluded that “*mitochondrial genomes are … shaped by somatic … selection throughout organismal lifetimes”*. Our analysis using an alternative approach, however, failed to detect any signs of selection dynamics in mtDNA mutations of the Serrano mutational dataset. Additional support for the idea of somatic selection was provided by Serrano’s assertion of the existence of “*extreme selective pressures impressed by nuclear–mitochondrial matching to re-introduce the ancestral allele”*. We have previously shown that these *‘reverse mutations’* are the result of sample-to-sample cross-contamination (Franco et al., 2024); thus, this aspect will not be discussed here.

## Results

### Somatic mutations are intermixed with germline mutations: a potential purifying selection bias

It is generally appreciated that germline mtDNA mutations are under purifying (germline) selection (Stewart et al., 2008), (Greaves et al., 2014) and therefore carry signatures of purifying selection. Thus, if somatic mutations happen to be intermixed with germline mutations, then the resulting composite mutational set would have a germline selection signature, and we would have erroneously concluded that the entire set is under purifying selection. We therefore asked whether the Serrano dataset might contain a significant admixture of germline mutations which could have affected the estimates of selection. An initial pilot inspection of the Serrano data showed that a proportion of mutations in the dataset were shared between animals of the same lineage, but not between unrelated animals of different lineages (Figure 1). This implied that such mutations were most likely originally generated in the germline, prior to the separation of the germ lineages leading to the different animals. Thus, they should have been subject to negative germline selection, and Serrano’s estimates of somatic mtDNA selection may be subject to bias. We therefore opted to reevaluate the presence of selection in Serrano data.

**Figure 1.**
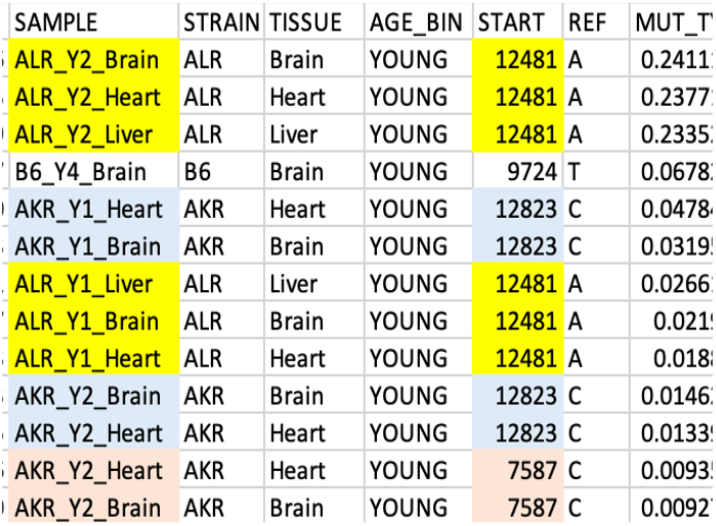
A screenshot of a portion of a worksheet that contains all synonymous mutations of Serrano et al. ranged by the decreasing mutant fraction. The mutations are colored by their sample of origin to show that a high proportion of synonymous mutations are likely germline mutations as they are shared between related animals.

### The alternative measure of selection: the ‘cumulative curve’ analysis

To assess selection, we opted NOT to use the traditional synonymity ratio metric, such as the hN/hS metric used by Serrano et al. The reason is that the suspected bias incurred by germline mutations admixture is expected to strongly depend on the mutant fraction (MF) because germline mutations are on average of higher fraction than somatic mutations, and thus the proportion of contaminating germline mutations and the incurred bias on selection estimation is expected increase at higher MF. This implies that for proper analysis, mutations need to be stratified/binned by MF. However, the binning structure itself is a source of bias, as convincingly shown by Serrano et al., who reported that selection appeared as predominantly negative or positive, depending on whether data was binned or not.

We thus used the ‘cumulative curve’ method originally proposed by Yuan et al. (Yuan et al., 2020) instead of the synonymity ratio. This approach essentially evaluates the pace of the fundamental progression of somatic mutations from nascent to high mutant fractions, and, eventually, to homoplasmy (Coller et al., 2001). This progression is driven by random genetic drift and, in some cases, selection. A faster progression results in a higher proportional mutational load being comprised by mutations of high mutant fraction (i.e., more clonally expanded) and thus is exactly what cumulative curve detects. To quantify and visually present this distribution of mutational load, the fraction of each given mutation is depicted as a function of the cumulative proportion of the of the number of mutations with fractions equal or higher than the fraction of the given one (**Figure 2**).

**Figure 2.**
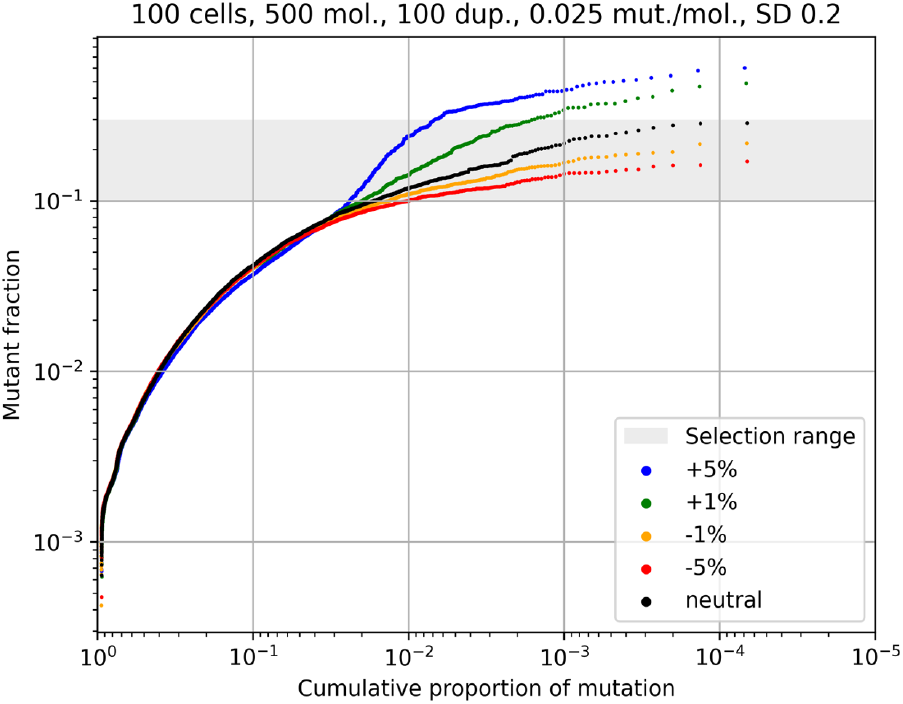
The “Cumulative curve” analysis of simulated data demonstrates the effectiveness of detection of selection of somatic mtDNA mutations: The dynamics of a population of cells with 500 mtDNA per cell was simulated for 100 duplications either without selection (black), or with positive (+1% green and +5% - blue) and negative (−1% - orange, −5% – red) selection imposed in the range 0.1< MF <0.3 (shaded).

The advantage of this approach is that in a dynamic system where mutations are generated de novo and subject to random drift and selection (like somatic and germline mutations are), the mutation dynamics are represented as a continuous trajectory, where nascent (low MF) mutations are predominantly on the left and ‘older’ (high MF) mutations are on the right. Note that this convenient left-to-right progression of mutations is achieved by plotting cumulative fraction in reverse. We use a logarithmic scale to emphasize ratios rather than absolute differences and to fairly cover wide dynamic ranges. In this representation, the local slope of the trajectory represents, in relative terms, the average rate at which mutations progress towards high fraction and homoplasmy, specifically at the corresponding MF. This, in turn, allows one to compare the rates, for example, of synonymous and nonsynonymous mutations and therefore estimate selection. This can be done continuously at any given MF, which makes the potentially biased binning of data by MF unnecessary. Most importantly for our discussion, mutations under selection and those that are not lie on different, typically well-separable, trajectories due to the difference in rate of their accumulation.

Figure 2. illustrates the above principles using simulated data (see Methods). Simulated mutations subject to the neutral drift only (which is expected of synonymous mutations) lie on a monotonous neutral drift trajectory (colored black in **Figure 2**). When the simulation includes positive selection, the mutations’ trajectory expectedly deviates positively around the mutant fraction (the Y-axis) of the selection onset (0.1 in this example) and returns on the original slope upon cessation of selection (MF=0.3). As seen in the figure, the magnitude of this effect qualitatively reflects the intensity of selection (compare green and blue trajectories). Conversely, negative selection causes the trajectory to deflect negatively from the neutral trajectory (orange and red trajectories). This example demonstrates the utility of cumulative curves for the assessment of selection. For synonymity analysis, it’s important that if the pace of the progression is equal for synonymous and nonsynonymous mutations, then there is no net selection. If, however, the nonsynonymous mutations’ trajectory diverges from the synonymous trajectory, that implies the presence of selection in the direction of the deviation.

### An admixture of germline mutations causes the illusion of purifying selection

To reassess somatic selection in the Serrano mutational dataset, we constructed cumulative curves separately for all synonymous and nonsynonymous mutations extracted from the dataset (**Methods**). **Fig.3A** shows the results of this analysis: synonymous (green) and nonsynonymous (red), with a best matching neutral drift simulation (blue) added for reference. The initial impression of **Fig.3A** is puzzling. At face value, this looks like strong purifying selection: the trajectory of non-synonymous mutations falls much further below the synonymous trajectory-as if the progression of nonsynonymous mutations to higher MFs is suppressed by purifying selection. Moreover, we confirmed (not shown) that mutations in this range bear clear negative selection signature (an excess of synonymous mutations), which seemingly confirms the purifying selection idea.

**Figure 3.**
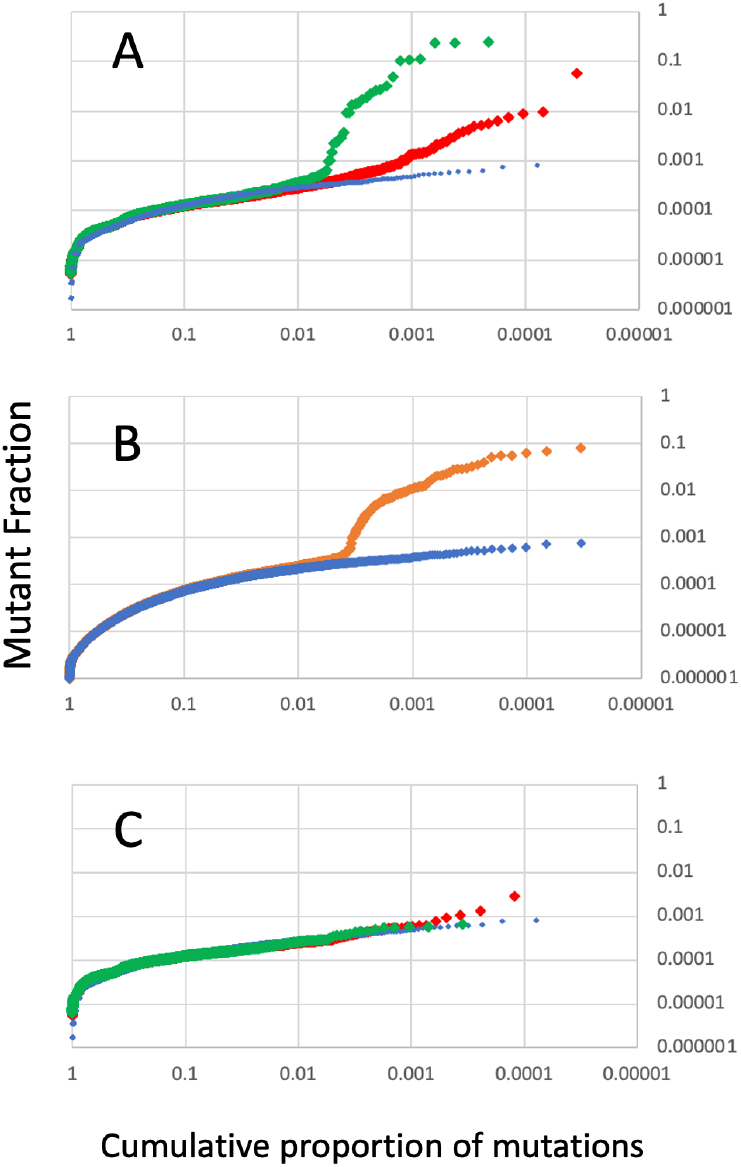
Cumulative curve comparison of synonymous (green) and nonsynonymous (red) mutations of the Serrano et al dataset, with a best matching neutral drift simulation (blue) for reference. **A**: all mutations. **B**: A qualitative simulation demonstrating the effect of an admixture of ‘simulated germline mutations’ to simulated somatic mutations shaped by neutral genetic drift (see Methods for details of the simulation). **C**: Cumulative curve of the Serrano

The ‘red flag’ for this interpretation, however, is that if this were purifying selection, one would expect the nonsynonymous mutations’ trajectory to digress negatively from the neutrally simulated values, as depicted in the negatively selected simulations in **Fig.2**, (orange and red curves). Instead, they diverge positively from this neutral trend, which should also represent the experimental synonymous mutations. Instead, the synonymous trajectory sharply deflects upwards from both nonsynonymous and neutral trajectory as if synonymous mutations are under positive selection, which is if course, impossible, and thus poses a conundrum that requires resolution.

To address this problem, we reasoned that because cumulative curves are merely a way to present the mutant frequency distribution, an upward deflection of the synonymous curve around MF∼0,001 in Fig3A may not imply acute somatic selection but reflect an excess of large synonymous mutations with MF>0.001, as compared to neutral drift model expectation, present for reason other than selection. Because we have shown above that Serrano somatic mutations indeed are mixed up with germline ones (**Fig. 1**), an obvious candidate for the source of such an excess of mutations is the admixture of germline mutations. Under this hypothesis, the synonymous trajectory (green) runs higher that nonsynonymous one (red) merely because germline mutations that are abundant in that MF range are highly synonymous (the proportion of synonymous germline mutations is very high), which is a well-known result of purifying germline selection (Stewart et al., 2008). The transition’s position at MF∼0.001 also makes sense, because germline mutations primarily originate from primordial germ cells (PGC), which contain about 1,000 mtDNA (Cao et al., 2007). In other words, the selection-like pattern in **Fig.3A**. is not a result of acute somatic selection, but of past time germline selection. In support of this hypothesis, we demonstrated, in a pilot simulation (**Fig. 3B**), that an admixture of simulated ‘germline mutations’ to simulated mutations driven by random drift (See **Methods** for details) results in an upwards deflection of the cumulative curve highly reminiscent of that observed in the Serrano data (**Fig.3A**).

To directly test the hypothesis of the germline admixture, we removed germline mutations from the Serrano dataset and repeated the cumulative curve analysis. For this purpose, germline mutations were inclusively defined as mutations shared between animals of the same mtDNA lineage (see **Method**s for details). In full agreement with the prediction, the removal of germline mutations fully abolished the dramatic upwards deflection of synonymous and nonsynonymous mutations (**Figure 3C)**. This confirms that the selection-like effect seen in Fig.3A is merely a result of germline mutation admixture. Moreover, this implies that mutations located on the ascending segments of the cumulative curves in Fig 3A are enriched with germline mutations, which indeed proved to be the case (>90% of them are shared between related animals’ germline mutations).

We noted that the prevalence of synonymous mutations among high-fraction mutations might be alternatively explained (as suggested by Serrano et al) by purifying selection acutely taking place in somatic tissues, but being limited to high-fraction mutations, only, e.g. because of a frequency threshold. To test this plausible hypothesis, we compared the cumulative curve pattern in young and old animals. The pattern is essentially identical (**Supplementary Figure 1**), implying that purifying selection did not take place during the lifespan, though we can’t exclude that it took place earlier than 4 months, i.e., the age of the young mice in this experiment.

### Overall, somatic mutations are not under selection

The removal of germline mutations naturally allows to assess the true somatic selection potentially masked by the overwhelming germline selection. **Fig. 3C** shows cumulative curve analysis of the dataset with germline mutations removed. Remarkably, the curves for synonymous, nonsynonymous and simulated neutral mutations are essentially indistinguishable from each other. This implies that *overall*, somatic mutations are *not* under selection. This, however, does not necessarily mean that there is no somatic selection of mtDNA mutations whatsoever. Fairly rare cases of selection, and/or positive and negative selection cases that are balanced on average, will not affect the shape of the curve because it is an aggregate characteristic of very large number of mutations. In fact, examples of somatic selection are widely known, as discussed in the Introduction. **Fig.3C**. does imply, however, that Serrano et al. data do not support an overwhelmingly positive or negative selection of somatic mtDNA overall, within which is in accord with a previous report (Sanchez-Contreras et al., 2023).

Of note, in **Fig. 3C**, there is a mild (merely 4 mutations) upturn of the non-synonymous curve (red) at MF>0.001 relative to the neutral simulation curve (blue) and synonymous mutations curve (green). This excess of nonsynonymous mutations hints of potential positive selection at higher mutant fractions. Although there is not enough data to conclude on significance of this trend, this observation is interesting, particularly because it echoes the observation by (Li et al., 2015) of positive selection at higher mutant fractions (MF>0.005). More research is needed to meaningfully explore this intriguing connection.

Mitochondria are known to change with individual’s age, so, to exclude the possibility that we might have missed selection because the data were not stratified by age, we additionally constructed cumulative curves separately for young and old animals (**Supplementary Figure 2, A and B**). The two sets of cumulative curves are very similar, implying that significant age-dependent differences are unlikely.

Finally, we asked whether the absence of systematic overall selection of somatic mutations is a general observation. We therefore repeated our analysis on a different dataset, e.g. the somatic mutation dataset that was generated with the same methodology (though at lower sequencing coverage) and published earlier by the Kennedy laboratory (Sanchez-Contreras et al., 2023). Unfortunately, Kennedy’s dataset lacks a reliable information on the kin relationship among experimental animals. We therefore defined somatic mutations as those which appeared exactly once in the entire dataset (this is a stricter criterion than that used in Fig. 3C, so for this comparative analysis in **Supplementary Figure 3**, Serrano dataset was subject to the same, stricter, criterion. Remarkably, the cumulative curves for the two datasets are highly similar, which increases our confidence in the conclusions.

### There is no evidence of subunit-specific selection

Whereas our analyses indicated that there is no overall selection on somatic mutations in the entire mitochondrial genome, there is a possibility that individual mitochondrial genes are under gene-specific positive and/or negative selection. Such cases may mutually compensate each other leading to the perception of the global absence of selection in the whole genome. To this end, a possibility of subunit-specific selection was implied by Serrano et al, who performed a detailed analysis of potential signatures of selection among the 13 individual protein coding genes in mtDNA for all tissue types and ages, resulting in over 250 comparison graphs. This analysis led them to conclude that some of the subunits were under selection, and ND5 was singled out as most consistently demonstrating a negative selection signature. Because of concerns over the unusually large number of comparisons which constitute multiple independent hypotheses and relative sparsity of the data as it gets divided in so many subsets, we decided to reevaluate these conclusions. We focused on ND5 as the most likely candidate for negative selection and constructed cumulative curves for its mutations (**Fig. 4**). As seen in the figure, the curves match precisely the whole genome curves in **Figure 3C** (except they are, as expected, shifted to the left, because of the smaller total number of mutations), implying that there is no indication of gene-specific selection of the ND5 mutations.

**Figure 4.**
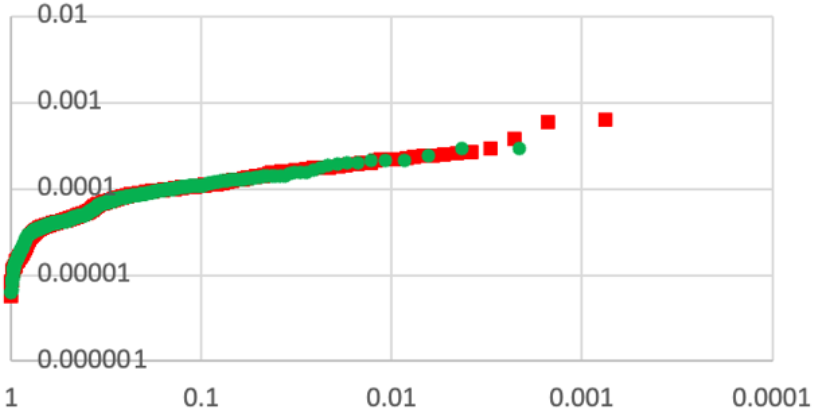
Cumulative curves for the somatic mutations of the ND5 subunit gene.

## Conclusions

Our analysis demonstrates that previously reported signatures of purifying selection acting on somatic mitochondrial DNA (mtDNA) mutations are largely explained by the inadvertent inclusion of germline mutations, which are subject to negative selection and are more likely to be synonymous and of higher mutant fraction. By separating germline from somatic mutations and applying a cumulative curve approach to assess the dynamics of synonymous and nonsynonymous variants, we found no evidence of systematic selection acting on somatic mtDNA mutations across tissues or ages. This finding reconciles conflicting reports in the literature and suggests that while isolated cases of positive or negative selection may exist—particularly at high mutant fractions—there is no pervasive selection shaping the somatic mtDNA mutational landscape. Furthermore, we found no support for subunit-specific selection, including in ND5, which had been previously highlighted. These results emphasize the importance of distinguishing germline and somatic variants and caution against interpretation of aggregated mutational data when assessing selection in mitochondrial genomes.

## Methods

### Data sources and pre-processing

Specific mutation information was available as a supplemental table from the Serrano et. al. and (Sanchez-Contreras et al., 2023) papers. The Serrano dataset was additionally cleaned; we removed all ‘mutations’ that were identical to the polymorphic positions among the mouse mtDNA lines used in the study as we have previously demonstrated that these ‘mutations’ result from sample cross-contamination (Franco et al., 2024).

### Construction of cumulative curves

The cumulative curves used in this manuscript were adapted from Yuan et al. To construct these curves, the mutant fractions of the subset of interest is ranged inversely (largest to smallest). Then, the cumulative proportion of all mutations with mutant fraction equal or larger than that of the given mutation to the total mutant load is calculated for each mutation. The curve is generated by plotting the mutant fraction of each mutation as a function of the cumulative proportion of the number of mutations with a mutant fraction smaller than the given mutation (in reverse).Then, for better visibility of trends, each axis is log transformed. An additional convenience of this representation is that the area under the cumulative curve represents the relative cumulative mutant fraction of mutations with a fraction smaller than the Y-coordinate. So, although the logarithmic scale makes visual perception distorted, it greatly emphasizes the contributions of the large mutations.

### Classification of somatic and germline mutations

The mutations reported by Serrano et al. were measured in somatic tissues. However, some of these mutations might have originated from and existed in the germline and therefore could have been subject to germline selection, resulting in bias of our estimates of somatic selection. Because the main goal of this study is to reevaluate the selection of somatic mutations, it was crucial to ensure that the data is clean from non-somatic mutations. **Figure 5** below explains our strategy to determine the mutations’ origin.

**Figure 5.**
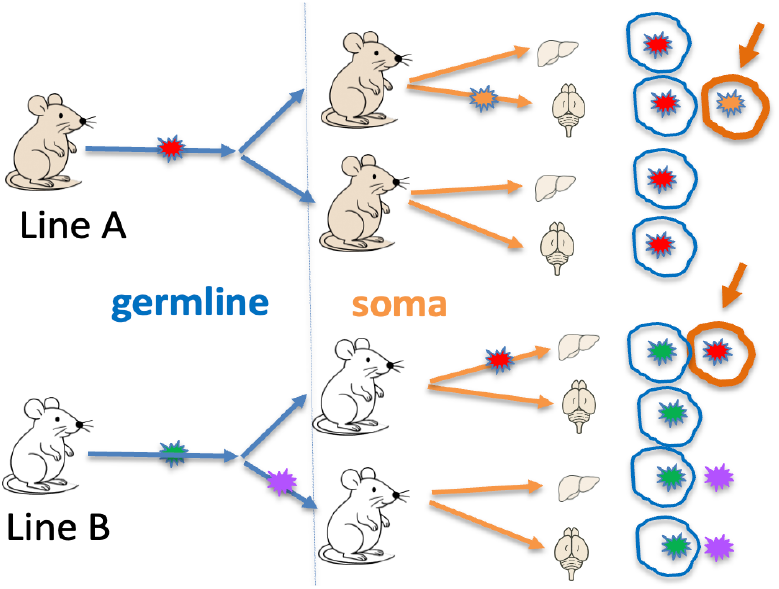
Definition of mutation origin. Somatic mutations (orange circles and arrows): unique to a somatic tissue sample among related animals (i.e. carrying the same mtDNA lineage). Can be present in unrelated animals. indicated by the orange circles. Germline mutations: shared among related animals (blue circles). Uncertain origin: mutations shared between tissues of an animal, but not among related animals – may be either germline or from early development (purple).

**Figure 6.**
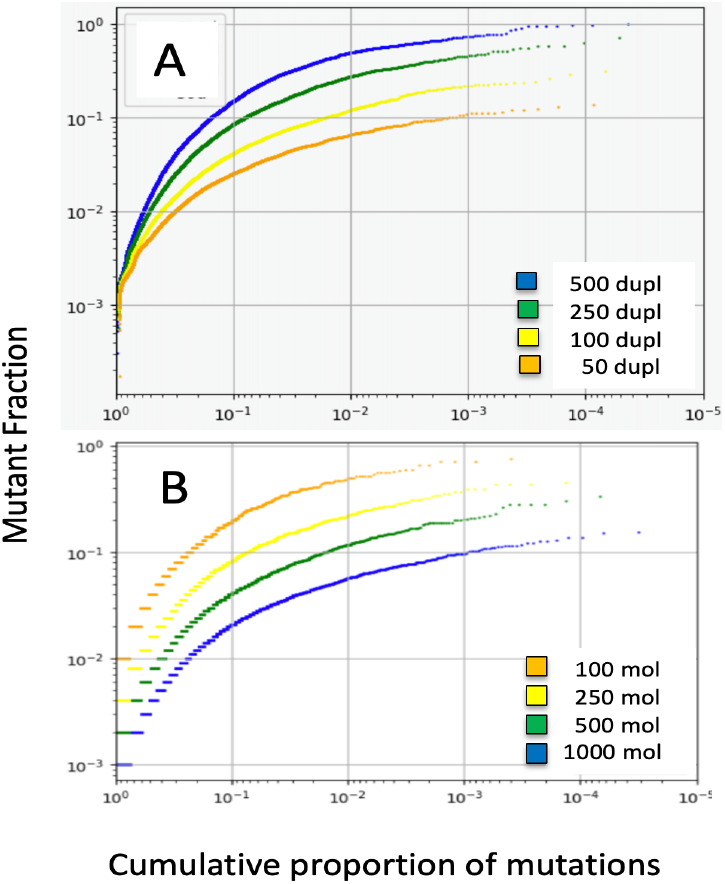
Parameters of simulations that affect the shape of cumulative curves: A – number of cell duplications B – number of mtDNA copies per cell. Color coding shown in the legends.

The classification of mutations was based on the understanding that mouse lineages carrying conplastic mtDNA were subject to 10 generations of backcrossing, which implies that all animals carrying the same mtDNA are highly interrelated and very distantly related to animals carrying different mtDNA lineages. Thus, any germline mutations are likely to be shared only between mice of the same mtDNA lineage. A mutation is deemed somatic if it is detected once in one tissue of an animal. If a mutation is detected in at least two tissues, it is likely that the mutations was generated (and should have accrued substantial presence) in the cellular populations that separated into the tissues before the separation, which makes the mutation a likely candidate for being present in the germline (purple mutation in the figure). We did allow, however, the same mutation to appear in an unrelated animal (red mutation), because such mutation must be a result of an unrelated mutational event and thus does not challenge somatic status of the mutation in question. Also, the red mutation in this example will be listed once as a somatic mutation (at mutant fraction that was recorded for this mutation in mouse #1 of line B. Mutations at the same genomic position will be also listed 4 times as germline mutations in brain and liver of both mice of line A, at the corresponding MFs.

We emphasize that this classification is not fully accurate, misclassifications are expected. For example, we may deem a mutation that showed in a single tissue as somatic, whereas in fact it was created in the germline and failed to show up in the second tissue because it was lost due to random drift. On the other hand, two identical mutations might have been created independently in related animals, but we would classify both mutations as germline. Even with these caveats in mind, this classification nevertheless definitely enriches both somatic and germline mutations. The effectiveness of this enrichment is empirically confirmed by the dramatic changes of the cumulative curves that we observe upon removal of mutations classified as germline (compare **Fig 3A and 3C**).

### Simulations of neutral intracellular mutation dynamics

In the simulation of the dynamics of neutral (synonymous) mutations, each simulation begins with a population of virtual cells with a given number of mitochondrial genomes. The cells accrue mitochondrial mutations at a given rate:, i.e., mutations are added onto a randomly chosen genomes in the cell at the beginning of a replication cycle. In each replication, the cells undergo duplication and random segregation of their mitochondrial genomes, which mimics the genetic drift. This is achieved by copying each molecule in the cell and randomly selecting the original genome number from the duplicated pool to assign to the daughter cell. In the interest of computational space, the genomes that would have been assigned to the alternate daughter cell are discarded. The populational variability that is lost in this caveat is regained in the number of total cells run in the simulation (which still allows for each cell’s duplications to remain efficient). At the end of the given number of replications, the mutation fractions of the present mutations are calculated as the number of times the mutation is present in the cell/the number of mitochondrial genomes in the cell, where the minimum value is 0 and maximum value is 1.

### Conversion of intracellular simulation results into mutant fractions in bulk tissue

#### The concept

The simulation described above reflects the intracellular dynamics of mutations. The actual MF measurements reported in Serrano et al. and in (Sanchez-Contreras et al., 2023) pertain to bulk tissue samples, so the simulated intracellular mutational fractions need to be ‘converted’ to be comparable to experimental measurements. The experimental mutations include both somatic and germline mutations. The ‘conversion’, however, is very different for these types of mutations since germline mutations span the entire body approximately evenly. For example, a 10% neutral mutation in a germ cell will result in approximately 10% mutation in all tissues of the pup that eventually grows from this germ cell. In contrast, a 10% somatic mutation in a somatic cell ends up as in a very small fraction of mtDNA of a bulk tissue sample. The actual fraction as measured for such a somatic mutation depends on a few potentially known and unknown factors. First, it depends on the size of the sample. More specifically, it depends on number of non-mutant mtDNA copies in the sample, because this number is in the denominator of mutant fraction. ‘Non-mutant’ mtDNA are molecules that do not carry the mutation in question, but they may carry other mutations. In other words, non-mutant mtDNA ‘dilutes’ the apparent MF. On the other hand, in most cases, cells with a given mutation are not alone in the bulk sample but are a part of a somatic clone of cells carrying the same mutation (such as a colon crypt (Greaves et al., 2014)). The size of mutant somatic clone increases mutant fraction, but, even with that, mutant fraction of a somatic mutation in a tissue sample typically is orders of magnitude lower than its fraction in the somatic cell from which this mutation has arisen (‘bulk tissue factor’). Likewise, the fractions of somatic mutations are typically orders of magnitude lower than those of germline mutations. Note that this factor does not apply to single cell somatic mutation studies, e.g. (Nekhaeva et al., 2002) or (Korotkevich et al., 2024). The complexity is further increased since, to make the complete dataset, dozens of samples are added together, and conclusions are based on aggregated data.

#### Somatic mutations

The above reasoning shows that there is no a priori way to make simulated mutations comparable to the experimentally observed ones because not enough information about internal architecture of the tissue is available. So, we have chosen an empirical approach: to qualitatively fit the neutral simulated cumulative curve to the experimentally observed neutral somatic cumulative curve (**Fig.3C, green**). The parameters of simulations that affect the shape of the curve are the number of molecules per virtual cell and the number of duplications (**Fig.6**). Additionally, the ‘bulk tissue factor’ moves the curve down parallel to itself, without altering its shape. We started with ‘common sense’ parameters of 1000 molecules per cell and 100 cell duplications as a reasonable expectation for an ‘average’ somatic cell. Of note, these two parameters are not independent as in genomic drift models an increase of the population size is compensated by an increase of the number of duplications. That is, the model would behave in the same way with 3000 molecules and 300 duplications (assuming the total number of mutations is kept the same for statistical stability reasons). The resulting cumulative curve has a shape essentially indistinguishable from the experimental curve, except, as expected, the MF were much higher. Then it was an easy task to determine the bulk tissue factor (which was determined to be ∼1/300) that qualitatively superimposes the simulated curve with the experimental one..

#### Germline mutations

As discussed above, simulated mutations can be used to represent germline mutations without ‘bulk tissue’ adjustment. As for a basic simulation parameters, we considered that the effective size of the mtDNA bottleneck is considered ∼ 50 (Arbeithuber et al., 2020). This is equivalent to the drift in cells with 1000 molecules for 20 duplications, which is about how many duplications germline cells go through per generation. Germline mutations that are observed in the Serrano et. al. dataset, especially the largest, most ‘influential’ ones, have likely passed several organismal generations (∼20 cell duplications in germline each), so we assumed it fair to use 1000 molecules, 100 duplications (i.e. the same parameters as for somatic mutations) to fairly emulate the dynamics of germline mutations. Of note, the simulations described here (e.g., shown in Fig 2B) are not intended for precise reproduction or exact fitting experimental data. Rather, it was intended as a qualitative illustration of the general effect on the shape of the cumulative curve.

#### Combined somatic/germline simulation (Fig.3B)

To correctly simulate the distribution of mutations in the bulk tissue and make it comparable experimentally observed mutations one needs to combine the somatic and the germline simulated mutations at the right ratio. The actual ratio of somatic and germline mutations is difficult to predict *a priory*, so we used the measurable parameter – the ratio of the total mutant load of somatic and germline mutations in the Serrano dataset as a guide. In Serrano dataset, this ratio is about 6. So, to make a combined somatic/germline mixture of mutations that fairly represents the Serrano’s dataset, we put together a set of ‘somatic mutations’, i.e. simulated mutations with frequencies divided by 300, and a set of germline mutations, i.e. mutations from the same simulation with non-adjusted fractions. The number of the added non-adjusted ‘germline’ mutations was such that their combined mutant fractions was 6-fold of the combined mutant fraction of the set of the ‘somatic’ mutations. The resulting curve is shown in **Fig.3B** (orange). It shows a pretty good visual fit to the experimental curve based on the Serrano dataset (**Fig.3A**, green).

#### Simulation of selection

Simulations that implement selection utilize the same framework as the neutral version but modify the probability of a mutation being transmitted to the daughter cell after duplication. The effects of selection are applied within specific mutant fraction ranges, with selection strength that is represented as a percentage, positive, or negative(e.g, 4% positive between 0.3-0.5 mutant fraction). During each round of replication, after any necessary mutations are added but before duplication and division occurs, the mutant fraction of each mutation in a cell is calculated. If a mutation has a mutant fraction that is within the range of selection, each molecule that contains the mutation has likelihood proportional to the strength of selection of being duplicated or removed from the cell at that time (before the cell-wide replication process) depending on the direction of selection. For example, if the selection parameters listed above are implemented and a cell with ten molecules contains a mutation with 0.4 mutant fraction, then each of the four molecules containing the mutation will independently have an *additional 4%* chance of being duplicated. More specifically, each of these four molecules will be assigned a random value between 0-1. If the value falls below 0.04 (4%), then an identical copy of that specific molecule will be added to the cell. This additional molecule will not independently undergo the selection process in the same round of replication. At the end of the selection process, all of the molecules in the cell including any added molecules are duplicated and the number of molecules that the cell should contain is randomly chosen to populate the daughter cell. If it were negative selection in this example, each of the four molecules would each have a 4% chance of being removed from the cell before the cell-wide replication process. This effectively changes the likelihood of the selected mutant to appear in the next consecutively daughter cell. At the end of each simulation, the result is a list of mutant fraction values that correspond to the prevalence of a given mutation in one cell. The simulations used in this manuscript were adapted into a webapp called MitoSim using Render and can be found at https://sim-app.onrender.com/.

## Supplement

**Supplementary Figure S1.**
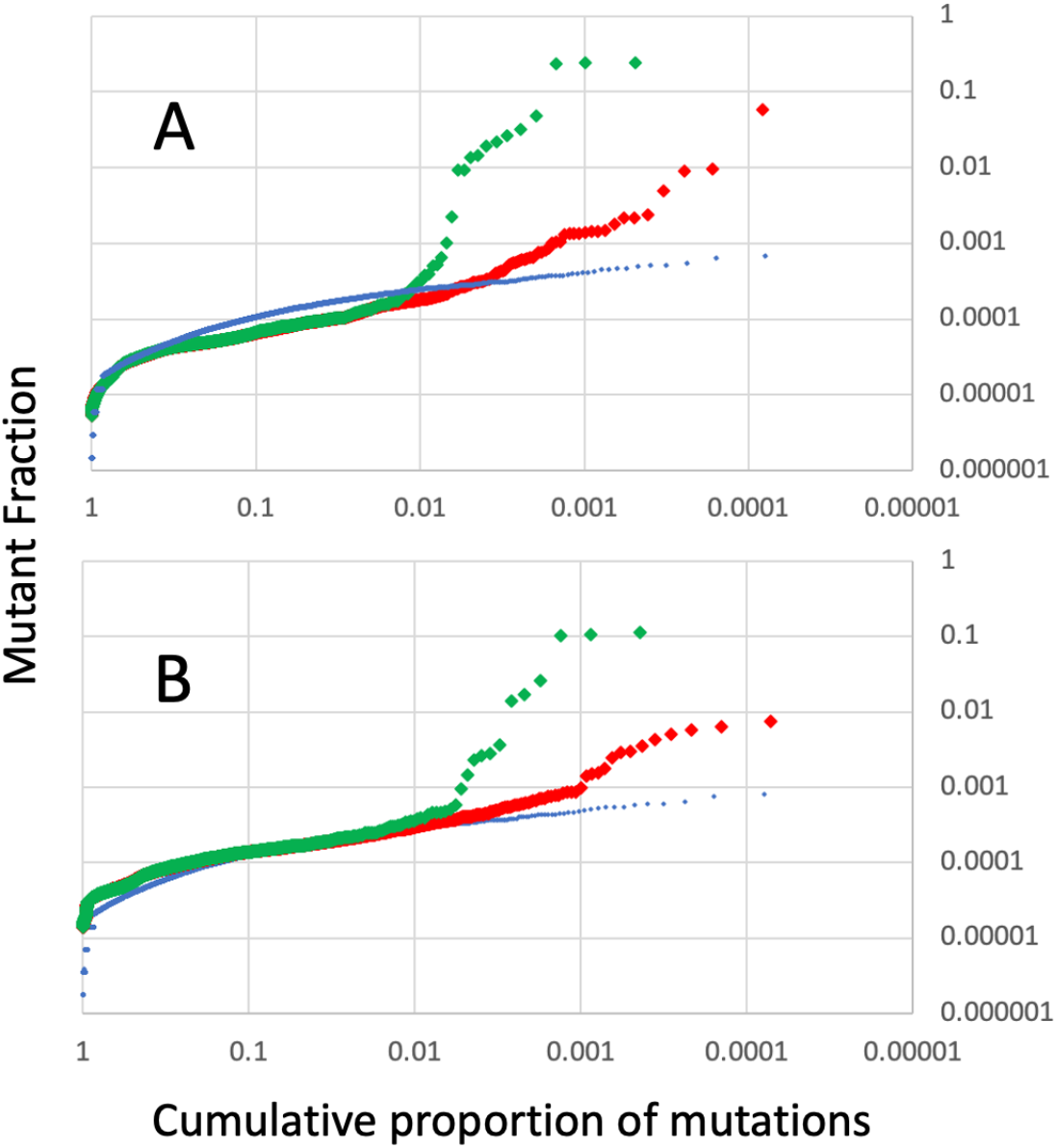
Germline mutations: Young vs. Old. Purifying selection of the germline mutations does not proceed in somatic tissues, at least during the lifespan (between young (top) and old (bottom)). Green – germline, red – somatic mutations, blue – neutral simulation (see Methods for details).

**Supplementary Figure S2.**
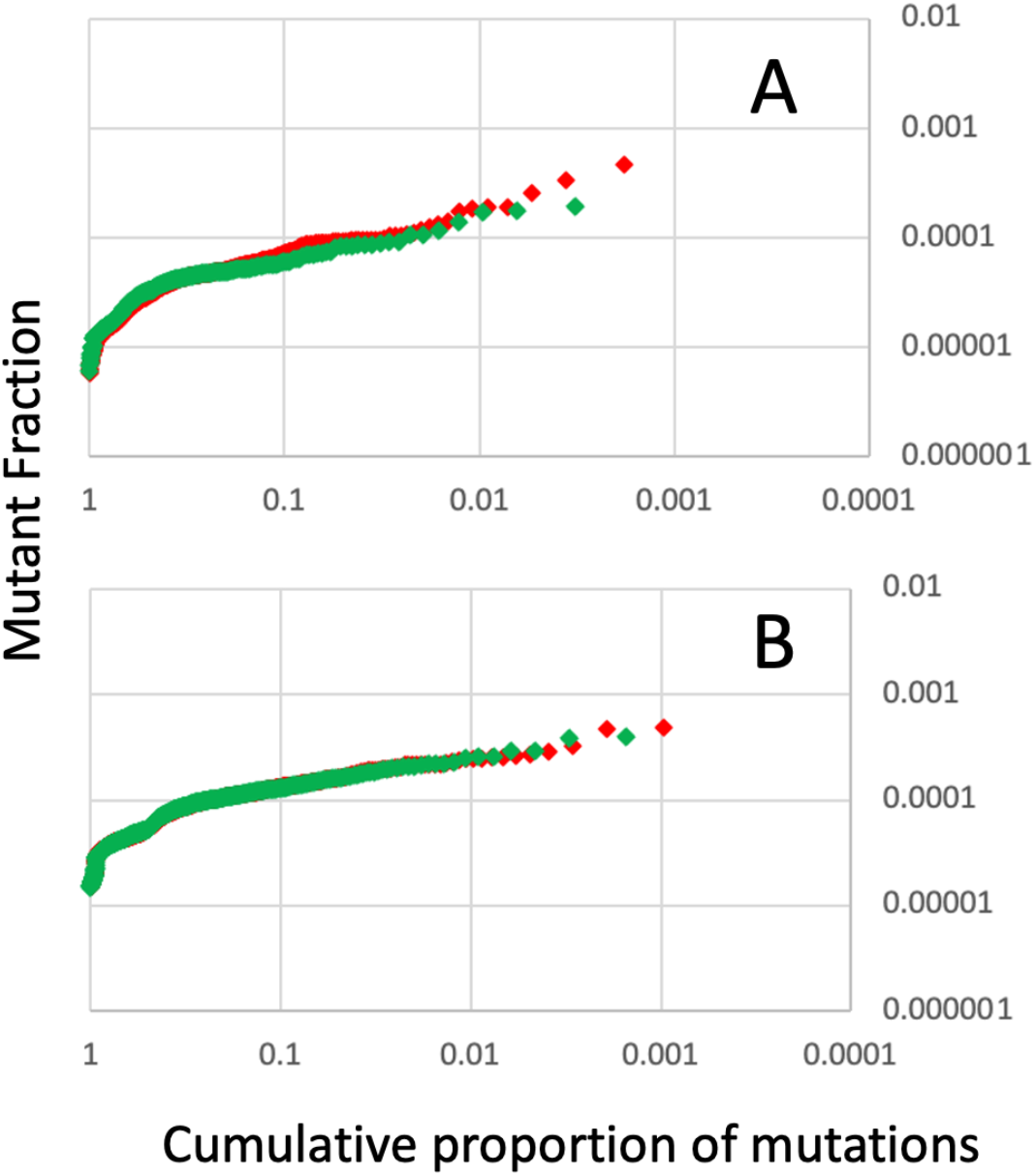
Somatic mutations: Young vs Old. Somatic mutations are not subject to selection either in young or old tissues and there is no lifetime-long trend of selection either. Cumulative curves are shown for somatic mutations of the young (A) and old (B) tissues. Green – germline, red – somatic mutations.

**Supplementary Figure S3.**
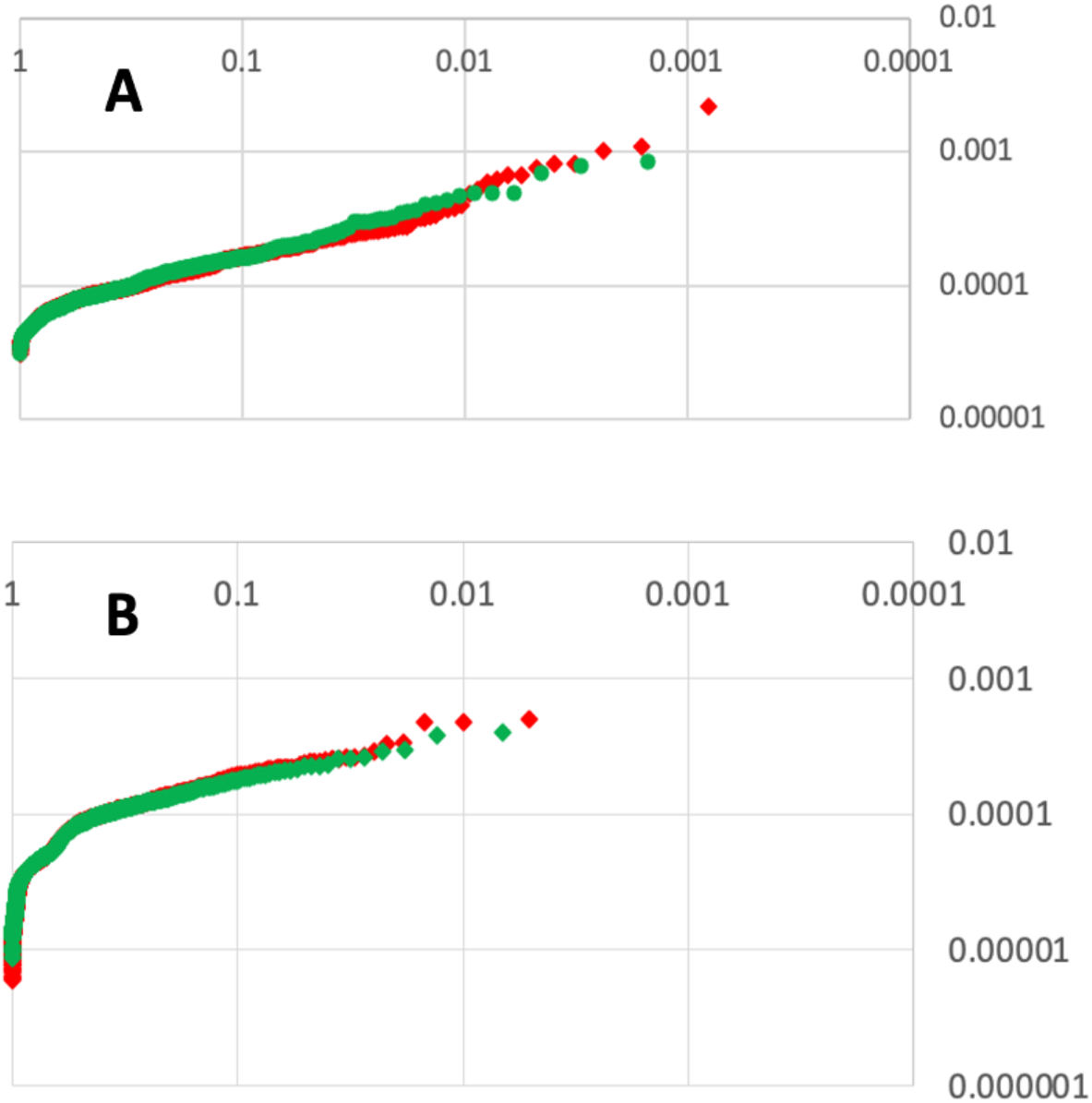
Serrano et al. vs. Sanchez-Contreras et. al. Cumulative curves for the somatic (single occurrence) mutations of the Sanchez-Contreras et. al 2023 dataset (**A**), compared to Serrano et al. dataset curve constructed exactly in the same way (and differently from Fig3C) (**B**). The difference in the dynamic range of the two graphs is due to a lower sequence read coverage in the Sanchez-Contreras et. al 2023 dataset. Green – germline, red – somatic mutations.

